# The Ies6 subunit is essential for INO80-mediated nucleosome organization

**DOI:** 10.1101/2025.11.21.689789

**Authors:** Ashish Kumar Singh, Felix Mueller-Planitz

**Affiliations:** Institute of Physiological Chemistry, Faculty of Medicine Carl Gustav Carus, Technische Universität Dresden, Fetscherstraße 74, 01307 Dresden, Germany; Molecular Biology, Biomedical Center, Faculty of Medicine, Ludwig-Maximilians-Universität München, 82152 Planegg-Martinsried, Germany; Epigenomics, Proliferation, and the Identity of Cells, Department of Developmental and Stem Cell Biology, Institut Pasteur, CNRS UMR3738, 75015 Paris, France

**Keywords:** INO80, Ies6, Nucleosome positioning and spacing, NRL, Nucleosome remodeler

## Abstract

ATP-dependent nucleosome remodelers of the INO80 family regulate chromatin by sliding, spacing and positioning nucleosomes. The INO80 remodeler is organized into structural modules that regulate its remodeling activity. Here, we investigate the role of the Ies6 subunit within the Arp5-Ies6 module towards nucleosome positioning and spacing in *Saccharomyces cerevisiae*. We show that Ies6 is critical for establishing genome-wide nucleosome organization. Deletion of *IES6* reduces nucleosome spacing by 3 bp and disrupts regular nucleosome arrays across most genes. Surprisingly, deletion of *IES6* is synthetically lethal with the deletion of *ISW2*, a remodeler from the ISWI family, indicating functional redundancy in nucleosome organization. Notably, INO80 binding directly predicts the role of Ies6 in INO80-mediated nucleosome organization, whereas changes in gene expression do not correlate with altered nucleosome spacing or array regularity. Together, our results highlight the essential role of the Ies6 subunit in INO80-mediated chromatin organization.

## INTRODUCTION

Genomic DNA is wrapped around nucleosomes, which affects all genomic processes involving DNA transactions ^1,2^. Nucleosomes attain a stereotypical architecture downstream of the promoter region where equally spaced nucleosomes, known as regular nucleosome arrays, are aligned next to the transcription start site (TSS). The distance between two adjacent nucleosomes in a regular nucleosome array is referred to as the nucleosome repeat length (NRL) ^3,4^. How cells establish and maintain this organization in the face of disruptive transcription and replication processes is an active area of research. ATP-dependent nucleosome remodeling complexes play pivotal role in this process ^5^.

These nucleosome remodeling complexes can evict, slide and space nucleosomes as well as exchange canonical histones with histone variants. They can be classified into four main families: SWI/SNF, ISWI, CHD and INO80 ^5^. The INO80 remodeler plays essential regulatory roles in multiple processes including transcription, replication, DNA damage response and double-strand break repair ^6–8^. Together with the ISWI and CHD families, INO80 has the ability to slide and equally space nucleosomes *in vitro* ^9–12^. In cells, INO80 positions the first nucleosome (+1 nucleosome) downstream of the TSS and can generate regular nucleosome arrays ^13,14^.

The *S. cerevisiae* INO80 remodeler is a 15-subunit complex, consisting of the Ino80 ATPase subunit and three major regulatory modules – the Nhp10, Arp8 and Arp5/Ies6 modules ^15,16^. The Nhp10 and Arp8 modules can associate with extranucleosomal DNA and are proposed to act as sensors of linker DNA length and regulators of the enzymatic activity of INO80 ^17–20^. Consistent with such a regulatory role, deletion of *ARP8* in *S. cerevisiae* resulted in a shortened NRL, implicating that the Arp8 module in regulating the nucleosome spacing activity of INO80. Despite its clear involvement in sensing linker length *in vitro* ^19^, deletion of *NHP10* however did not alter the NRL, suggesting that this module is dispensable for nucleosome spacing ^14^.

In contrast to Arp8 and Nhp10 modules, the Arp5-Ies6 module acts as a central core component of the INO80 remodeler (Fig. 1A). It binds to DNA and histone moieties of the nucleosome core particle. It also contacts the acidic patch on the H2A-H2B dimer ^21,22^, which acts as an allosteric activator of INO80 activity. As such, the Arp5-Ies6 module appears to play an important role in regulating ATP hydrolysis and nucleosome sliding by the INO80 complex, as demonstrated *in vitro* with mono-nucleosome substrates. A mutant INO80 complex lacking Arp5-Ies6 is still able to bind and slide nucleosomes, albeit with decreased efficiency ^16,23–26^.

**Fig. 1.**
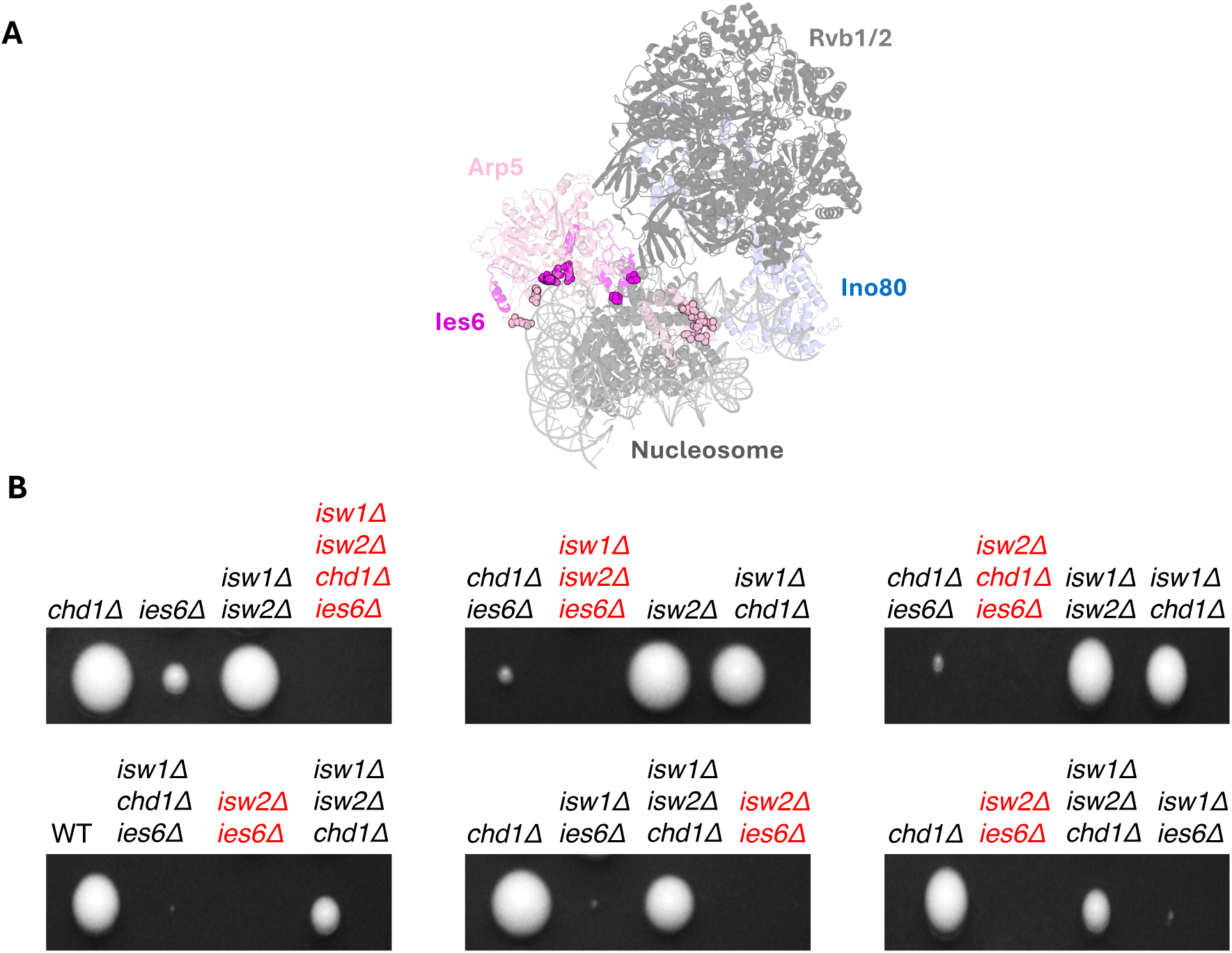
Deletion of *IESC* is synthetically lethal with deletion of ISW2, but not with deletion of *ISW1* or *CHD1*. (**A**) Structure of the *S. cerevisiae* INO80 complex bound to the nucleosome. Selected subunits are highlighted. Spheres denote residues from Ies6 (magenta) and Arp5 (light pink) that interact with the nucleosome. PDB code: 9C9G. (**B**) Representative tetrad dissection of diploids arising from TKO x *iesCE*. Tetrads were dissected on a YPAD plate, colonies grown for five days and replica-plated on minimal media lacking appropriate amino acids or on full media with cloNAT selection marker to identify genes lacking in the colonies. For additional results, see Supp. Fig. S1.

In yeast, deletion of either the Arp5 or Ies6 subunit is suggested to lead to a loss of the entire module from the INO80 complex ^27^. These mutants display disrupted yeast metabolic cycle and mitochondrial maintenance that parallel, but do not completely recapitulate the phenotypes observed upon deletion of the Ino80 ATPase subunit ^28,29^. Moreover, loss of the Ies6 or of the Ino80 subunits results in increased ploidy due to defective nucleosome organization near centromeres ^30^. At the transcriptome level, deletion of Arp5 or Ies6 alters the expression of more than 1,000 genes. Deletion of the Ino80 ATPase subunit produces an even more severe phenotype ^24,31^. These findings suggest that an INO80 complex lacking the Arp5–Ies6 module retain partial remodeling activity. Consistent with this, the Arp5–Ies6 module has been proposed to carry out both specialized roles within the INO80 complex and potentially independent functions outside of it in yeast and cancer cells ^28,32^.

Here, we test the role of Ies6 subunit in regulating nucleosome positioning and spacing in *S. cerevisiae* by analyzing the phenotypes associated with *IES6* deletion. Unexpectedly, we find that *iesCE* is synthetically lethal with *isw2Δ*, but not with *isw1Δ* and *chd1Δ*, suggesting that INO80 and ISW2 may regulate a similar subset of genes. Deletion of *IES6* decreased nucleosome positioning, NRL and regular arrays genome-wide. Surprisingly, changes in gene expression detected by RNA-seq upon deletion of *IES6* do not predict alterations to the chromatin architecture following the deletion, indicating that transcriptional alterations do not directly account for the chromatin defects observed in *iesCE* cells.

## RESULTS

### Deletion of IESC is synthetically lethal with deletion of ISW2, but not with deletion of ISW1 or CHD1

Nucleosome remodelers often act redundantly to position and space nucleosomes. We and others have previously shown that INO80 can contribute to nucleosome spacing within gene bodies of *S. cerevisiae*, in addition to Isw1 and Chd1 spacing remodelers ^13,14,25,33–36^. The role of the Arp5-Ies6 module in nucleosome organization by INO80 remains poorly understood. It forms multiple interactions with the nucleosome (Fig. 1A) and has been suggested to grip on to the nucleosome while the ATPase domain pumps DNA ^22^. Since *ARP5* deletion is lethal in W303 yeast strains ^37^, we focused on *IES6* in this study. *IES6* could be readily deleted through homologous recombination in W303 wild-type cells indicating that it is not an essential gene (for strain list, see Supp. Table 1). Consistent with previous results, deletion of *IES6* gave rise to slow-growing colonies ^30^ (Fig. 1B, Supp. Fig. S1).

Since the presence of other spacing remodelers (Isw1, Isw2 and Chd1) obscures INO80 activity *in vivo* ^14^, we attempted to delete *IES6* also in a triple knock out (TKO) strain carrying deletions of *CHD1* and both ISWI factors (*isw1E isw2E chd1E*) ^38^. Despite multiple attempts, we did not succeed in deleting the *IES6* gene in TKO cells via homologous recombination. Therefore, we hypothesized that deletion of *IES6* may be essential in the TKO but not WT cells, or in other words, that *IES6* shows synthetic lethality with one or more of the remodelers deleted in TKO cells.

To test for synthetic lethality of Ies6 with ISW1, ISW2 and Chd1 remodelers, we generated diploid cells by mating haploid strains lacking *IESC* (MATα) with the TKO strain (MATa) and performed tetrad dissection. If *IES6* becomes essential in absence of any of these remodelers, we would expect to recover no viable spores carrying the corresponding double, triple, or quadruple deletions. Indeed, no viable colonies were recovered with the quadruple deletion (*iesCE isw1E isw2E chd1E*) or with the double deletion of *iesCE isw2E* (Fig. 1B, Supp. Fig. S1; lethal combinations in red). In contrast, double deletions of *iesCE* with either *isw1Δ* or *chd1Δ* produced viable, albeit small colonies. Similarly, the triple deletion of *iesCE isw1E chd1E* also resulted in small colonies, whereas combinations such as *iesCE isw2E isw1E,* and *iesCE isw2E chd1E* were not recovered (Fig. 1B, Supp. Fig. S1). Taken together, loss of viability in the *iesCE* TKO strain is driven by a synthetic lethal relationship between *IES6* and *ISW2*, with *ISW1* and *CHD1* playing lesser roles.

### Ies6 is essential for INO80 nucleosome positioning and spacing activity

To investigate the role of Ies6 in nucleosome organization, we performed micrococcal nuclease sequencing (MNase-Seq) on WT and two independent *iesCE* colonies. Deletion of *IESC* led to a genome-wide decrease in nucleosome organization. Compared to WT cells, the +1-nucleosome and the downstream nucleosomes in the gene body were less well positioned as seen in gene-averaged composite plots. Furthermore, gene body nucleosomes were shifted upstream towards the TSS, suggesting that *iesCE* cells possess on average a shorter NRL (Fig. 2A).

**Fig. 2.**
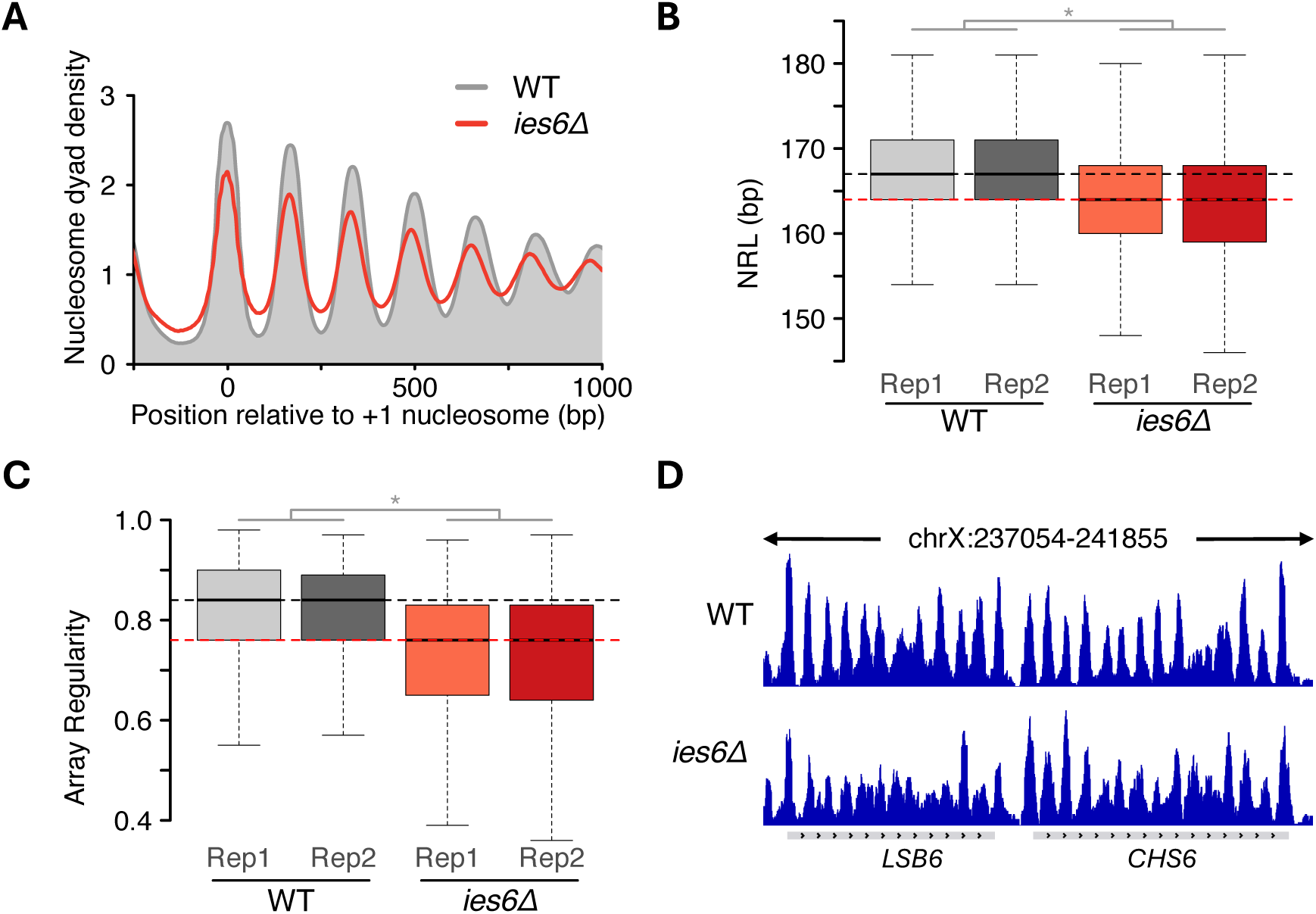
Deletion of *IESC* leads to genome-wide decrease in nucleosome positioning and spacing in *S. cerevisiae*. (**A**) Average nucleosome organization for all 5015 genes in the indicated yeast strains. (**B**) Boxplots showing NRL distribution in all yeast genes in WT and *iesCE* cells for two independent replicates, Rep1 and Rep2. (**C**) Same as (B), but for array regularity. Horizontal dotted lines indicate the average median values in WT (gray) and *iesCE* cells (red). (**D**) Genome browser snapshot over two representative genes in WT and *iesCE* cells.

To quantify changes to the structure of nucleosome arrays over each gene, we applied a previously published method for calculating NRL and array regularity ^14,34^. We found that the NRL decreased by 3 bp upon *IES6* deletion relative to WT (Fig. 2B). In addition, array regularity also declined, consistent with the reduced nucleosome organization observed in composite plots (Fig. 2C). These genome-wide trends were also evident at individual genes (Fig. 2D). Collectively, these results demonstrate that the Ies6 subunit is critical for establishing nucleosome organization *in vivo*.

We next asked to what extent *IES6* deletion phenocopies loss of the INO80 ATPase subunit. We previously showed that acute depletion of INO80 in W303 cells decreases the NRL by 3 bp ^14^, mirroring the decrease in NRL upon *IES6* deletion (Fig. 2B). Likewise, array regularity also showed a significant decrease upon depletion of the INO80 subunit (Supp. Fig. S2A).

To bolster these results that were obtained in the W303 yeast background, we inspected nucleosome array structure in BY4741 cells lacking the Ino80 ATPase using a published dataset ^25^. Consistent with results in the W303 background, deletion of Ino80 ATPase in BY4741 cells reduced NRL by 3 bp and decreased array regularity (Supp. Fig. S2B, C). Importantly, we observed a substantial overlap between genes showing decreased NRL in *iesCE* and in cells lacking the INO80 complex (Supp. Fig. S2D, E). Overall, these findings indicate that the INO80 remodeler regulates nucleosome organization in *S. cerevisiae* and that the Ies6 subunit plays an essential role within the INO80 complex, closely phenocopying the loss of the entire INO80 complex.

### Ino80 binding, rather than expression changes, predicts altered nucleosome organization following *IESC* deletion

To assess the causal basis of the effects observed upon *IESC* deletion, we compared INO80- bound genes with all other genes (Kubik et al. 2019). If Ies6 is important for INO80 activity, *IES6* deletion should produce a stronger disruption at INO80-bound genes than across the rest of the genome. Consistent with this expectation, INO80-bound genes showed a greater reduction in array regularity relative to non-bound genes following *IES6* deletion (Fig. 3A, B). INO80-bound genes also experienced a larger drop in the NRL (4 bp) compared to the remaining genes (2 bp; Fig. 3C).

**Fig. 3.**
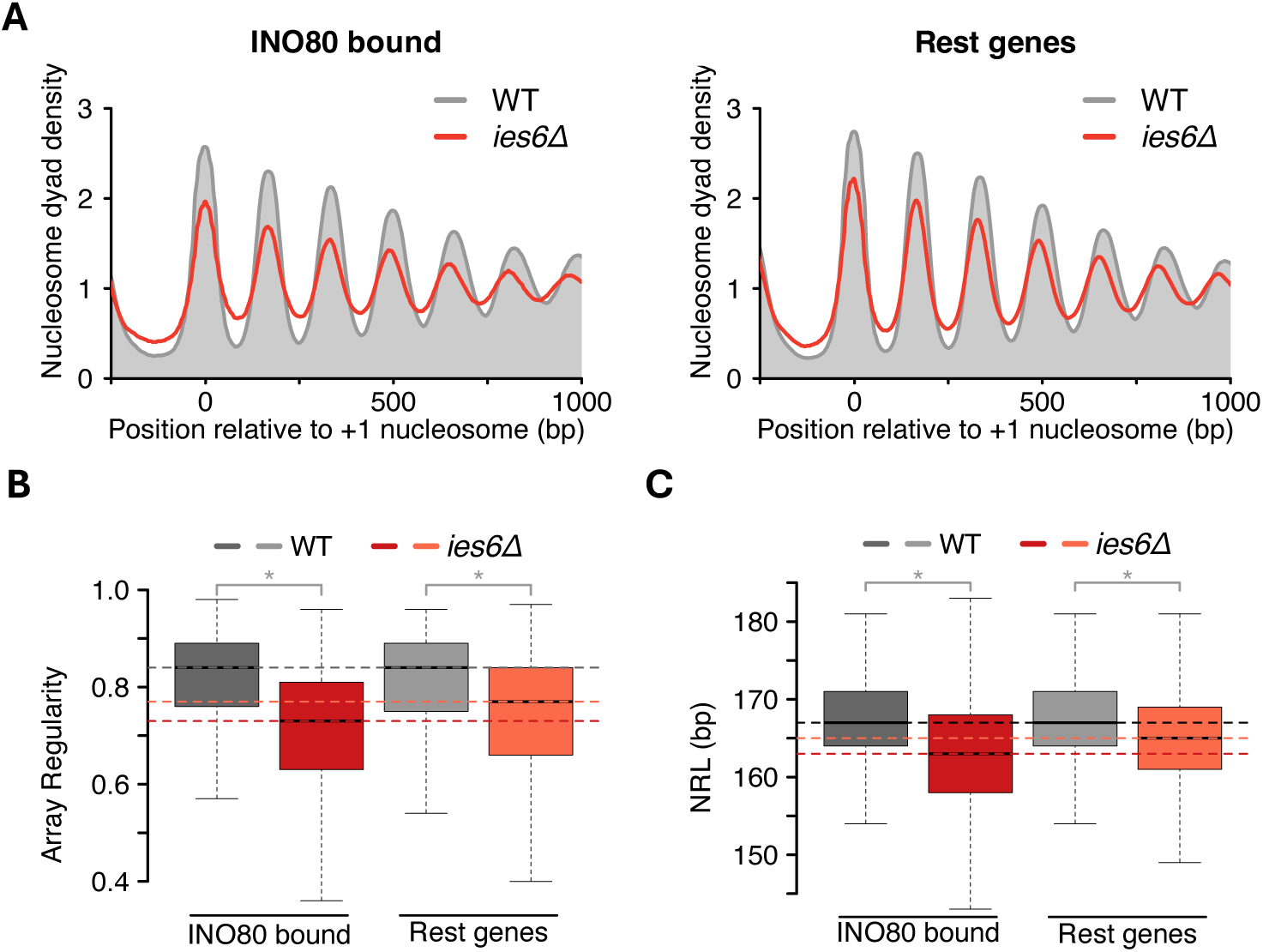
INO80-bound genes exhibit a greater reduction in nucleosome organization than other genes following *IES6* deletion. (**A**) Average nucleosome organization for all 1646 INO80 bound (left) and rest (right) of the genes in the indicated yeast strains. (**B**) Boxplots showing array regularity distribution in the same set of genes as in (A). (**C**) Boxplots showing NRL distribution in the same set of genes as in (A). Horizontal dotted line indicates the median NRL or array regularity in WT (gray) and *iesCE* (red) cells.

Analogous results were obtained when we tested the effect of the loss of the INO80 subunit: INO80 bound genes displayed a 4 bp decrease in NRL and a greater loss of array regularity compared to the rest of the genes (Supp. Fig. S3A, B, C) ^14^. These findings support a model in which INO80 induces even nucleosome spacing across the gene bodies it binds, with Ies6 serving as a pivotal subunit in this activity.

Motivated by these results, we next tested for a potential relationship between gene expression changes and changes in nucleosome organization over gene bodies upon Ies6 or Ino80 ablation. To test for such a relationship, we used published RNA-Seq datasets for *iesCE* and *ino80Δ* cells ^24,31^ and compared nucleosome features between genes with altered expression and the rest of the genome. Surprisingly, no preferential effects were observed. Following deletion of *IESC*, both differentially expressed and non-differentially expressed genes showed a similar decrease in NRL and array regularity (Fig. 4A-C, Supp. Fig. S4A, B). Similarly, genes with altered expression in *ino80Δ* cells did not exhibit preferential changes in nucleosome organization upon *IES6* deletion (Fig. 4D-F, S4C, D). Lastly, we examined nucleosome organization in *ino80Δ* BY4741 cells and again observed no preferential differences between genes with altered expression upon either *IES6* or *INO80* deletion (Supp. Fig. S5A-F). These results suggest that gene expression changes are not reliable indicators of altered nucleosome organization over gene bodies in *iesCE* or *ino80Δ* cells. Or, put differently, that an altered array structure over gene bodies in the deletion mutants is not predictive of changes in gene expression levels.

**Fig. 4.**
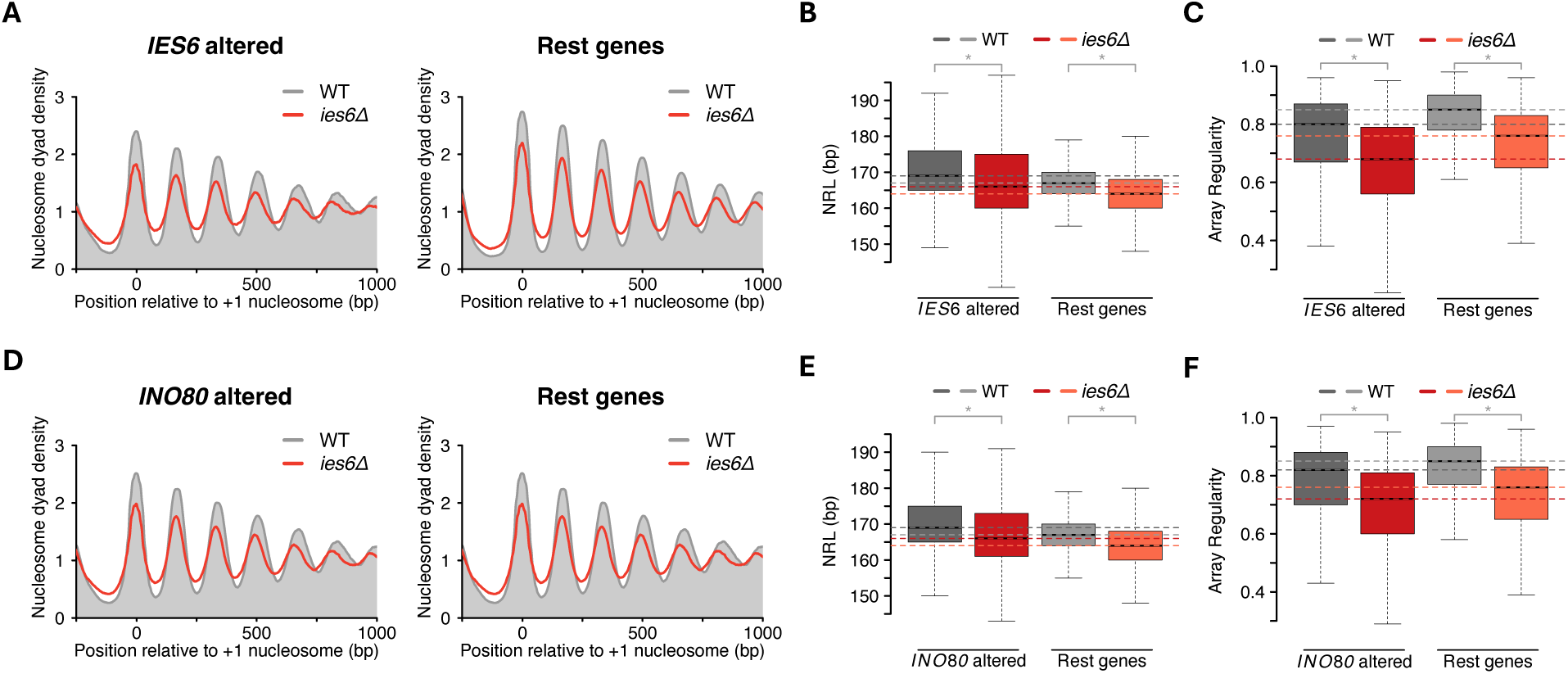
Gene expression changes upon *IES6* deletion do not correlate with changes in nucleosome architecture. (**A**) Average nucleosome organization in WT and *iesCE* cells for all 1304 genes with at least 1.5-fold change in gene expression in *iesCE* cells (left) and remaining genes (right). List of differentially expressed genes was obtained from ^31^. (**B**) Boxplots showing the NRL distribution in the same set of genes as in (A). (**C**) Boxplots showing array regularity distribution in the same set of genes as in (A). (**D**) Average nucleosome organization in WT and *iesCE* cells for all 1519 genes with at least 1.5-fold change in gene expression in *ino80Δ* cells (left) and for the remaining genes (right). List of differentially expressed genes was obtained from ^31^. (**E**) Boxplots showing the NRL distribution in the same set of genes as in (D). (**F**) Boxplots showing array regularity distribution in the same set of genes as in (D). Horizontal dotted line indicates the median NRL or array regularity in WT (gray shades) and *iesCE* (red shades) cells.

To further examine the relationship between the structure of nucleosome arrays and gene expression, we next compared genes with altered NRL to those with altered gene expression in *iesCE* and *ino80Δ* cells. Notably, we observed only a small overlap between genes showing NRL changes and those with differential expression as measured by RNA-Seq for both mutants (Fig. 5A, B). Even a stringent cutoff (>5 bp change in NRL) did not improve the overlap between the two gene sets (Supp. Fig. S6A, B). Taken together, these findings suggest that the limited overlap between NRL changes and RNA-seq data explains the lack of a direct effect of *IESC* or *INO80* deletion on nucleosome organization.

**Fig. 5.**
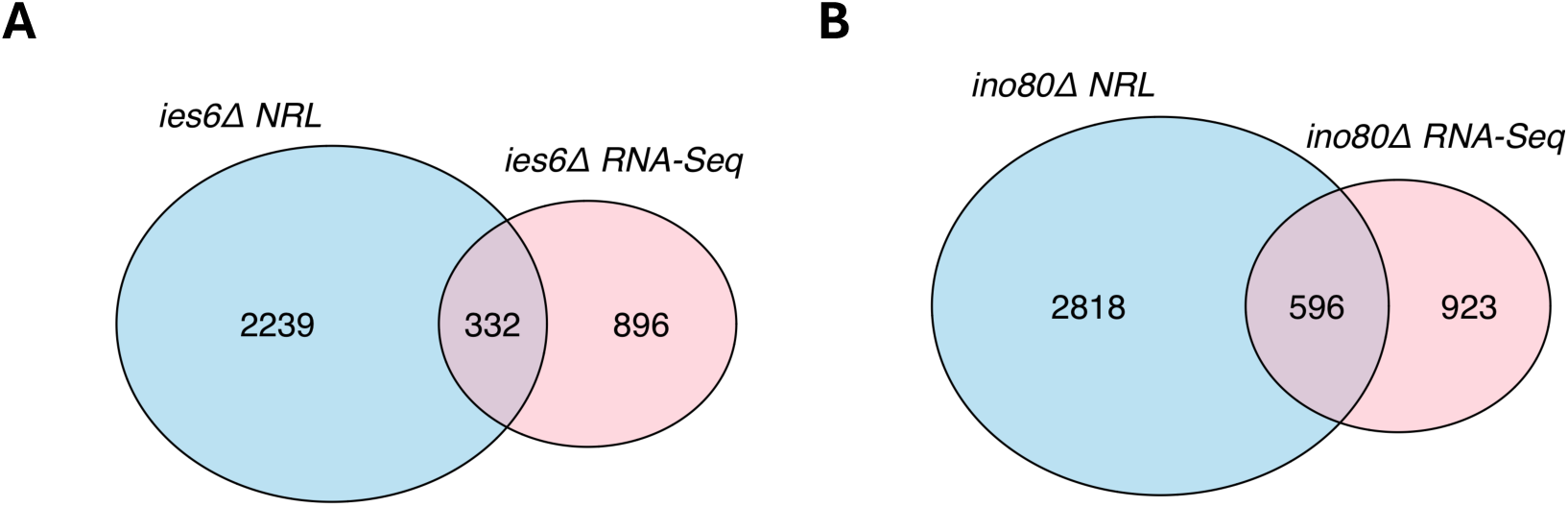
Genes with altered expression and NRL in *IES6* and *INO80* lacking cells show a partial overlap. (**A**) Venn diagram showing overlap of genes with altered RNA levels and NRL in *iesCE* cells. (**B**) Venn diagram showing overlap of genes with altered RNA levels and NRL in *ino80Δ* cells.

## DISCUSSION

Our study uncovers an essential role for the Ies6 subunit of the INO80 remodeler in establishing nucleosome organization across the *S. cerevisiae* genome. We show that the loss of Ies6 decreases nucleosome spacing and disrupts regular nucleosome arrays to a similar extent as deletion of the Ino80 ATPase. Our results strongly implicate Ies6 as a critical determinant of INO80-mediated chromatin organization *in vivo*.

Ies6 forms a structural module with Arp5. Together, these subunits establish multiple contacts with both nucleosomal DNA and histones (Fig. 1A), potentially gripping the nucleosome while the ATPase subunit translocates DNA ^22^. Loss of Ies6 likely destabilizes Arp5’s engagement with the nucleosome, reducing the efficiency of nucleosome sliding. In addition, Ies6 deficiency may impair Arp5’s ability to recognize the acidic patch of the H2A– H2B dimer, a critical allosteric activator of INO80 activity ^20^.

An unexpected outcome of our genetic analysis is the synthetic lethality observed between *ies6*Δ and *isw2*Δ. Whereas INO80 broadly shapes nucleosome architecture across the genome, Isw2 functions as a highly specialized remodeler that influences only a few hundred genes ^13,14,33,39^. Both remodelers are known to reposition +1 nucleosomes upstream into the nucleosome-depleted region, and we speculate that this shared activity underlies the synthetic lethality when both remodelers are absent. Consistent with this interpretation, growth defects have been reported upon simultaneous depletion of the Isw2 and Ino80 ATPases ^13^. In addition, both ISW2 and INO80 play critical roles in nucleosome organization near replication origins and within ribosomal DNA ^40,41^. Thus, the synthetic lethality between *ies6*Δ and *isw2*Δ may arise, at least in part, from combined defects in these genomic regions.

In addition to the lethal genetic interaction with ISW2, we also observed growth defects when *iesCE* was combined with *isw1Δ* or *chd1Δ*. Although these double and triple mutants remained viable, they displayed markedly slower growth than *iesCE* alone, suggesting partial redundancy between INO80 and other spacing remodelers in maintaining nucleosome organization. These results are also consistent with the model that multiple remodeler families collaborate to generate and preserve regular nucleosome arrays, yet certain remodelers play non-redundant roles at specific chromatin sites.

Another unexpected outcome of our study is the weak correlation between alterations in nucleosome organization and steady-state gene expression following deletion of either Ies6 or the Ino80 ATPase. This disconnect suggests that disruptions in nucleosome organization within gene bodies do not necessarily translate into measurable changes in steady-state transcription. Instead, more subtle architectural features, such as shifts in the positioning of the +1 nucleosome or reorganization within the nucleosome-depleted region upstream of it, may underlie the transcriptional alterations observed upon loss of Ino80 function. Furthermore, indirect or secondary consequences of Ino80 ablation, including perturbations in transcription factor recruitment, histone variant exchange, or an activated stress-response, could also contribute to the observed transcriptional outcomes. If not directly regulating gene expression, we speculate that the nucleosome-spacing activity of Ino80 serves to suppress cryptic transcription within intergenic regions and gene bodies, just like other spacing remodelers from the ISWI and Chd1 families ^14,31,42–44^.

## METHODS

### Yeast strain generation

All yeast strains used in this study (Supp. Table 1) were derived from the W303 background. To delete the *IES6* gene via homologous recombination, oligonucleotides oFMP1110 (gaaggttaaaattgtcatcatcatcagcgtgagaaagtcgaaacagatccccgggttaattaagg) and oFMP1111 (gaaaggttgtctacaagctaaaatacatacatacatatacaatgcgaattcgagctcgtttaaactgg) were used to amplify natMX6 marker from plasmid pFA6a-natMX6 (yFMP524). PCR products were purified from agarose gel and transformed into WT yeast strain, generating yFMP627 and yFMP628 lacking *IES6*. Gene deletion was confirmed by PCR using oligos oFMP1112 (cgatgacgacgactacct) and oFMP1113 (caaagtggagacgatgctg).

### Sporulation and tetrad dissection

The haploid strain lacking *IES6* (yFMP628) was mated with TKO (YTT227). Resulting diploids (yFMP468, yFMP469) were grown in YPA + 4% glucose media. Sporulation was induced in media lacking adenine, leucine and trpytophan (1% KOAc supplemented with uracil 5 mg/L and histidine 5 mg/L) at 23 °C for 4-5 days. Tetrads were dissected using a Singer MSM 400 dissection microscope onto YPAD plates and incubated at 30 °C for 5 days. Tetrads were replica-plated onto minimal media lacking appropriate amino acids or on YPAD plates with cloNAT selection and grown for an additional 3 days.

### MNase-Seq

Yeast nuclei preparation, MNase digestion and DNA isolation was performed as described previously ^14,45^. Briefly, cells were grown to OD600 0.8 in 500 ml YPAD media. Spheroplasts were prepared by treating cells with 2 mg zymolyase at 30 °C for 20 min, followed by washing with ice-cold 1 M sorbitol. Nuclei were resuspended in MNase digestion buffer (15 mM Tris– Cl pH 7.5, 50 mM NaCl, 1.4 mM CaCl2, 0.2 mM EGTA pH 8.0, 0.2 mM EDTA pH 8.0, 5 mM β-mercaptoethanol) and digested with increasing amounts of MNase for 20 min at 37 °C. DNA was purified via Proteinase K treatment, followed by Phenol: Chloroform: Isoamyl alcohol extraction and RNase digestion. Samples with ∼70% mononucleosome content were selected for library preparation. Sequencing libraries were prepared using the NEBNext Ultra II DNA Library Prep kit for Illumina and sequenced in 50 bp paired-end mode, yielding five to ten million reads per sample.

### Data analyses

MNase-Seq data were analyzed as described previously ^14^. Briefly, fastq files were mapped using Bowtie2 v2.2.9 to R64-1–1 sacCer3 genome with settings -X 500, -no-discordant, -no-mixed options ^46^. Fragments of 140-160 bp were selected and used for further. Nucleosome maps were generated by extending dyads to 50 bp. NRL and array regularity were calculated after subsampling all samples to 5 million reads using scripts (https://github.com/musikutiv/tsTools) which adapts previously published method ^34^. MNase-Seq data of cells lacking Ino80 ATPase in BY4741 background ^25^ were processed similarly as in this study. Genes were altered expression in *iesCE* and *ino80Δ* cells (fold change >1.5) were downloaded from GSE52000 and GSE77257 ^24,31^. Boxplots represent the median, interquartile range and 1.5X of the interquartile range. Statistical significance comes from a two-tailed Welch’s t-test performed on the median NRL values of two biological replicates. Pymol 3.0 was for structural visualizations. Residues interacting with the nucleosome were identified by selecting atoms from relevant subunits located within 4.0 Å of the nucleosome. The corresponding residues were then mapped and displayed as spheres to highlight potential contact sites.

## ACKNOWLEDGEMENTS

The authors thank Sigurd Braun and the Physiological Chemistry department at LMU Munich for providing access to the tetrad dissection microscope. We are thankful to David Clark for sharing scripts used to calculate NRL, to Tobias Straub and Tamas Schauer for general bioinformatics support and to Madhura Khare for help in tetrad dissection. We thank Philipp Korber and his lab for initial help with yeast genetics, Stefan Krebs and Helmut Blum at the Laboratory for Functional Genome Analysis (LAFUGA) for sequencing MNase-Seq libraries, and David Shore for providing a list of INO80-bound genes.

## AUTHOR CONTRIBUTIONS

Ashish Kumar Singh: Conceptualization, Data curation, Formal analysis, Investigation, Methodology, Validation, Visualization, Writing–original draft, Writing–review and editing. Felix Mueller-Planitz: Conceptualization, Formal analysis, Project administration, Funding acquisition, Supervision, Writing–review and editing.

## FUNDING

The work was supported by grants from the Deutsche Forschungsgemeinschaft (DFG project number 497659230 and SFB1064-A07). A.K.S. acknowledges financial support through a DAAD predoctoral scholarship.

## AVAILABILITY OF MATERIALS AND DATA

All high-throughput sequencing data have been deposited at the Gene Expression Omnibus under the accession number.

## ETHICS APPROVAL AND CONSENT TO PARTICIPATE

Not applicable.

## COMPETING INTERESTS

The authors declare no competing interests.

## SUPPLEMENTARY INFORMATION

**Supp. Fig. S1.**
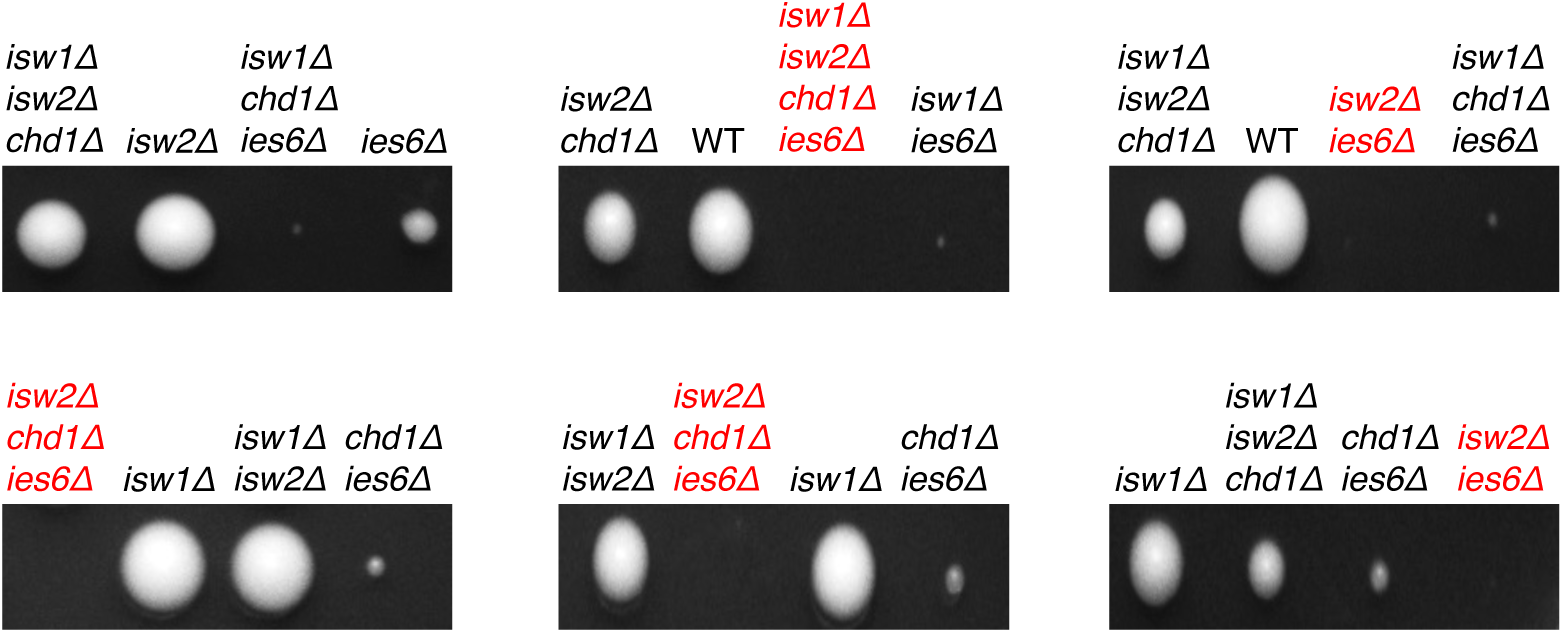
Further examples of tetrad dissection of diploids arising from TKO x *iesCE*. Tetrads were dissected on a YPAD plate, colonies grown for five days and replica-plated on minimal media lacking appropriate amino acids or on full media with antibiotic selection to identify genes lacking in the colony.

**Supp. Fig. S2.**
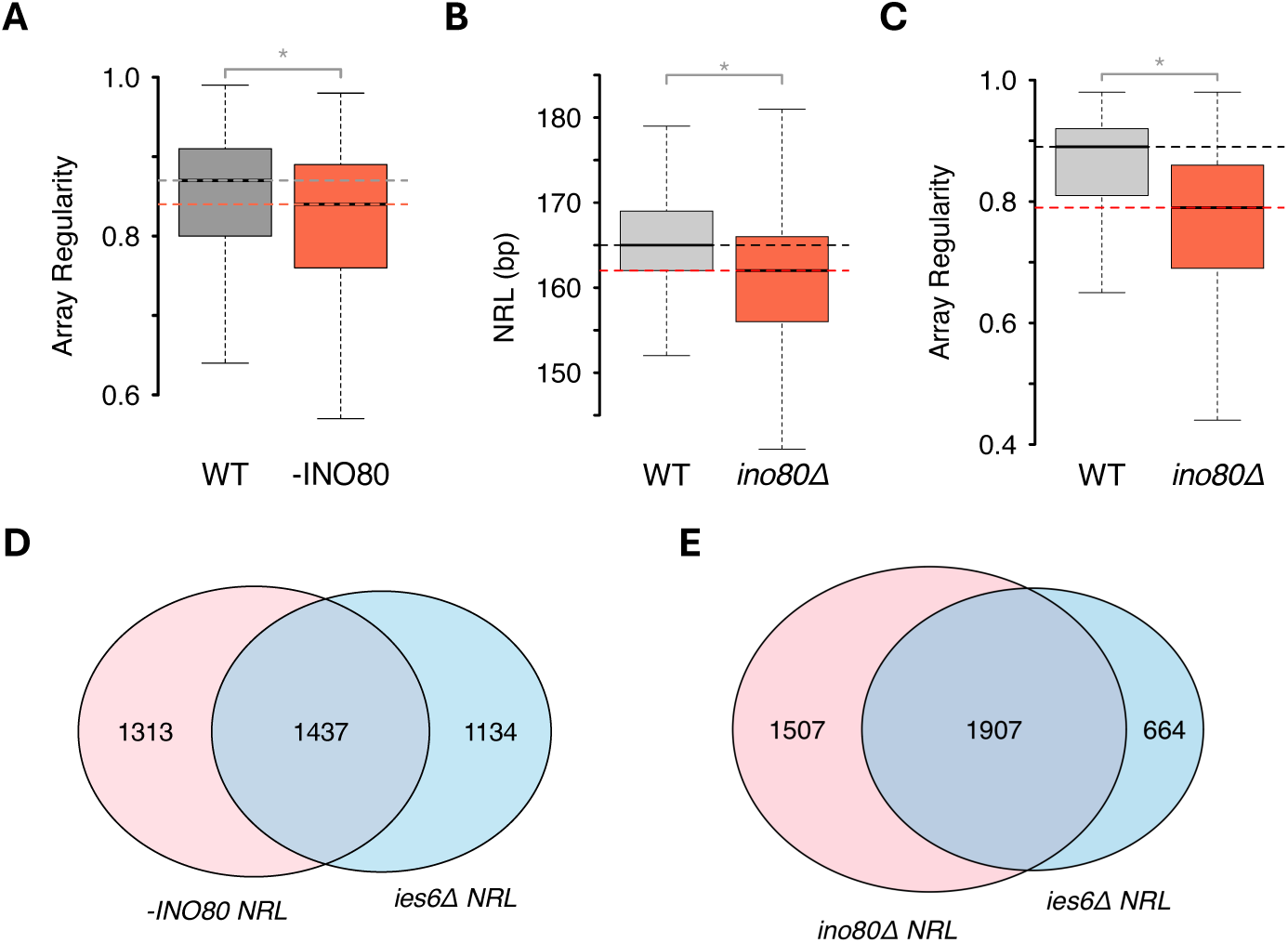
Deletion of Ino80 ATPase leads to lower NRL and array regularity in *S. cerevisiae*. (**A**) Boxplots showing array regularity distribution in 5015 yeast genes in WT and W303 cells depleted of INO80 via anchor-away method ^14^. (**B**) Boxplots showing NRL distribution in 5015 yeast genes in WT and BY4741 cells lacking Ino80 ATPase. (**C**) Same as B, but for array regularity. Horizontal dotted line indicates the median NRL or array regularity in WT (gray) and cells lacking INO80 (red shades) cells. (**D**) Venn diagram showing overlap of genes with altered NRL in *iesCE* and cells depleted of INO80 in W303 background ^14^. (**E**) Venn diagram showing overlap of genes with altered NRL in *iesCE* and *ino80Δ* cells ^25^.

**Supp. Fig. S3.**
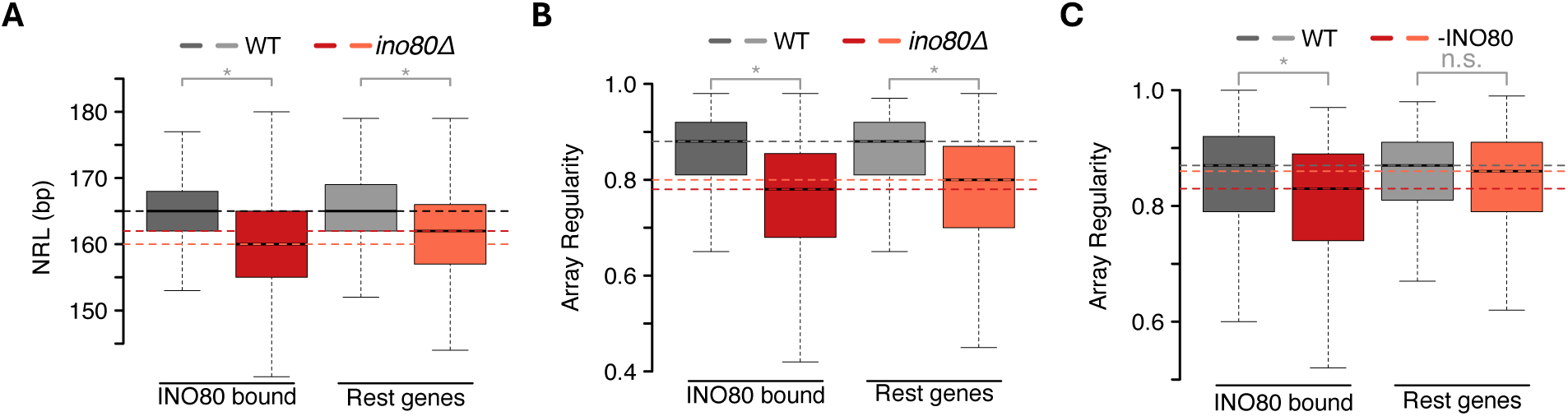
INO80-bound genes exhibit a greater reduction in nucleosome organization than other genes following ablation of Ino80. (**A**) Boxplots showing NRL distribution in 1646 INO80 bound (dark shades) and rest (light shades) in WT and *ino80Δ* cells ^25^. (**B**) Same as (A), but for array regularity. (**C**) Same as (A), but for array regularity distribution obtained after INO80 depletion ^14^. Horizontal dotted line indicates the median NRL or array regularity in WT (gray) and INO80 lacking cells (red shades).

**Supp. Fig. S4.**
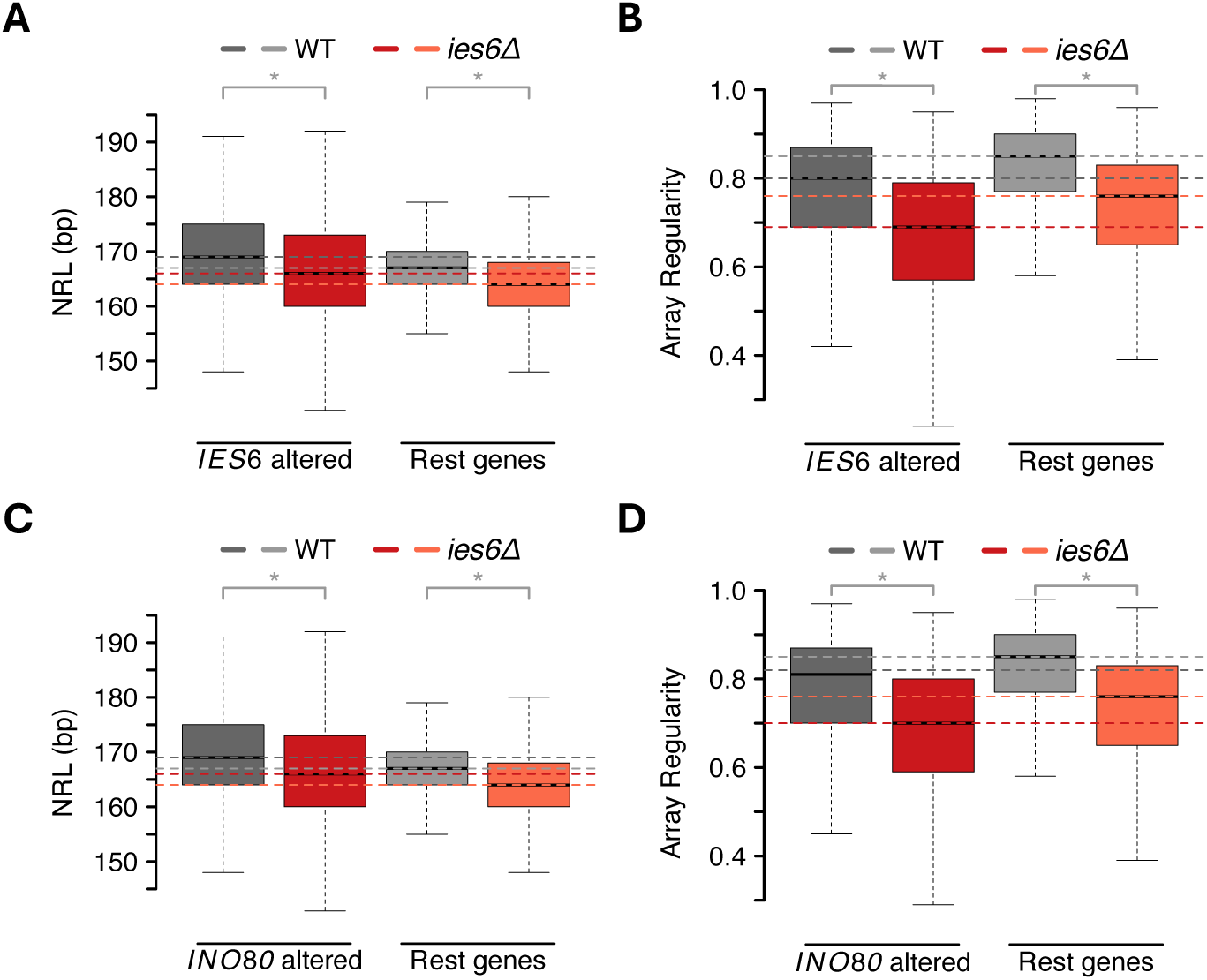
Genes with altered expression do not show different INO80 activity compared to unaltered genes in *iesCE* cells. (**A**) Boxplots showing NRL distribution in *iesCE* cells for all 1239 genes with at least 1.5-fold change in gene expression in *iesCE* cells (left) and remaining genes (right) in the indicated yeast strains. List of differentially expressed genes was obtained from ^24^. (**B**) Boxplots showing array regularity distribution in the same set of genes as in (A). (**C**) Boxplots showing NRL distribution in *iesCE* cells for all 1349 genes with at least 1.5-fold change in gene expression in *ino80Δ* cells (left) and rest (right) of the genes in the indicated yeast strains. (**D**) Boxplots showing array regularity distribution in the same set of genes as in (C). Horizontal dotted line indicates the median NRL or array regularity in WT (gray shades) and *iesCE* (red shades) cells.

**Supp. Fig. S5.**
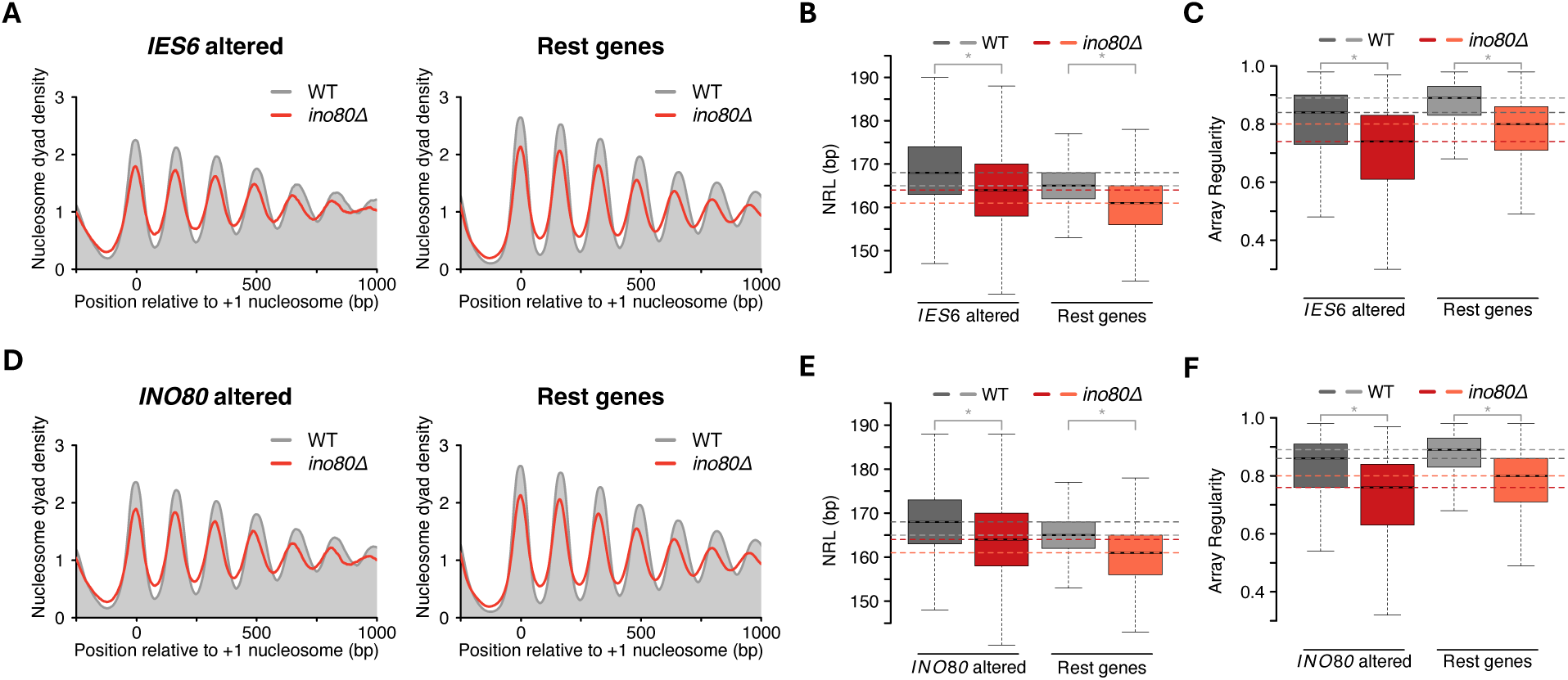
Genes with altered expression do not show different INO80 activity compared to unaltered genes in *ino80Δ* cells. (**A**) Average nucleosome organization in *ino80Δ* cells for all 1304 genes with at least 1.5-fold change in gene expression in *iesCE* cells (left) and rest (right) of the genes in the indicated yeast strains. List of differentially expressed genes was obtained from ^31^. (**B**) Boxplots showing NRL distribution in the same set of genes as in (A). (**C**) Boxplots showing array regularity distribution in the same set of genes as in (A). (**D**) Average nucleosome organization in *ino80Δ* cells for all 1519 genes with at least 1.5-fold change in gene expression in *ino80Δ* cells (left) and rest (right) of the genes in the indicated yeast strains. List of differentially expressed genes was obtained from ^31^. (**E**) Boxplots showing NRL distribution in the same set of genes as in (D). (**F**) Boxplots showing array regularity distribution in the same set of genes as in (D). Horizontal dotted line indicates the median NRL or array regularity in WT (gray shades) and *ino80Δ* (red shades) cells.

**Supp. Fig. S6.**
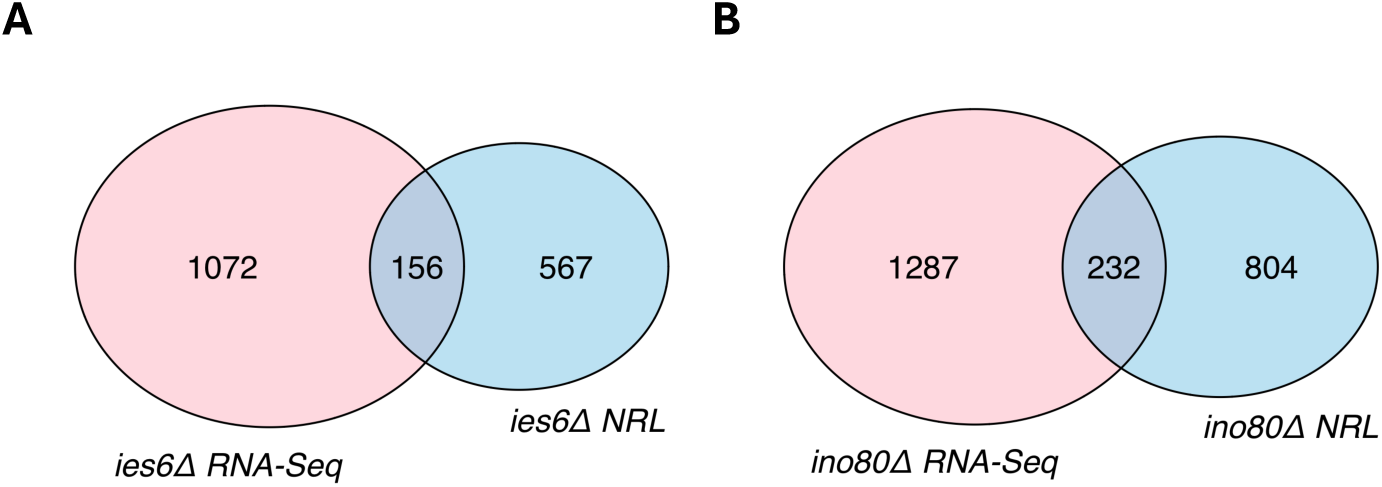
Genes with altered expression and more than 5 bp altered NRL in *IES6* and *INO80* lacking cells show a partial overlap. (**A**) Venn diagram showing overlap of genes with altered RNA levels and >5 bp change in NRL in *iesCE* cells. (**B**) Venn diagram showing overlap of genes with altered RNA levels and >5 bp change in NRL in *ino80Δ* cells.

**SUPPLEMENTARY TABLE 1:** List of yeast strains

| **Strain** | **Genotype** |
| --- | --- |
| W1588-4C<br>(yFMP013) | *MATa ade2-1 his3-11,15 leu2-3,112 trp1-1 ura3-1 can1-100 RAD5+* |
| YTT227<br>(yFMP014) | *MATa ade2-1 his3-11,15 leu2-3,112 trp1-1 ura3-1 can1-100 RAD5+ isw1Δ:ADE2 isw2Δ::LEU2 chd1Δ::TRP1* |
| yFMP627 | *MATa ade2-1 his3-11,15 leu2-3,112 trp1-1 ura3-1 can1-100 RAD5+ ies6Δ::NATMX6* |
| yFMP628 | *MATalpha ade2-1 his3-11,15 leu2-3,112 trp1-1 ura3-1 can1-100 RAD5+ ies6Δ::NATMX6* |
| yFMP468 | *MATa/MATalpha ade2-1/ade2-1 his3-11,15/his3-11,15 leu2-3,112/leu2-3,112 trp1-1/trp1-1 ura3-1/ura3-1 can1-100/can1-100 RAD5+/RAD5+ IES6/ies6Δ::NATMX6 C1* |
| yFMP469 | *MATa/MATalpha ade2-1/ade2-1 his3-11,15/his3-11,15 leu2-3,112/leu2-3,112 trp1-1/trp1-1 ura3-1/ura3-1 can1-100/can1-100 RAD5+/RAD5+ IES6/ies6Δ::NATMX6 C2* |

## Notes

### Competing Interest Statement

The authors have declared no competing interest.

